# Sensory Over-Responsivity: Parent Report, Direct Assessment Measures, and Neural Architecture

**DOI:** 10.1101/355628

**Authors:** Teresa Tavassoli, Anne Brandes-Aitken, Robyn Chu, Lisa Porter, Sarah Schoen, Lucy Jane Miller, Molly Rae Gerdes, Julia Owen, Pratik Mukherjee, Elysa J. Marco

## Abstract

**Background:** Sensory processing differences are common across neurodevelopmental disorders. Thus, reliable measures are needed to understand biologic underpinnings of these differences. This study aims to define a scoring methodology specific to tactile (TOR) and auditory (AOR) over-responsivity. Second, using MRI Diffusion Tensor Imaging, we seek to determine whether children with AOR show measurable differences in their white matter integrity.

**Methods:** This study includes children with AOR and TOR from a mixed neurodevelopmental disorders cohort including autism and sensory processing dysfunction (n= 176) as well as neurotypical children (n= 128). We established cut-off scores for over-responsivity using the parent report: Short Sensory Profile (SSP), and the direct assessment: Sensory Processing-Three Dimensions:Assessment (SP-3D:A). Group comparisons, based on AOR phenotype, were then conducted comparing the white matter fractional anisotropy in 23 regions of interest.

**Results:** Using the direct assessment, 31% of the children with neurodevelopmental disorders had AOR and 27% had TOR. The Inter-test-agreement between SSP and SP-3D:A for AOR was 65% and TOR was 50%. Children with AOR had three white matter tracts showing decreased fractional anisotropy relative to children without AOR.

**Conclusions:** This study identified cut scores for AOR and TOR using the SSP parent report and SP-3D:A observation. A combination of questionnaire and direct observation measures should be used in clinical and research settings. The SSP parent report and SP-3D:A direct observation ratings overlapped moderately for sensory related behaviors. Based on these initial structural neuroimaging results, we suggest a putative neural network may contribute to AOR.

## Background

Sensory processing dysfunction (SPD), manifest as difficulty interpreting the sensory world in an adaptive way, is common across children with neurodevelopmental disorders (NDD), including children who meet the categorical label of autism spectrum disorders (ASD) [1], [2]. Under the umbrella of SPD, there are three primary subtypes: difficulties modulating sensory input, difficulties discriminating sensory information, and difficulties with sensory-based motor control [3], [4]. While these challenges can exist independently, they often co-occur. The Diagnostic and Statistical Manual-5 (DSM-5), now includes hyper-or hypo-reactivity to sensory input (characteristic of sensory modulation) as a core criteria for ASD which has prompted additional interest and focus on sensory modulation [5].

Previous research suggests that one aspect of sensory modulation, sensory over-responsivity (SOR), occurs most frequently in the auditory and tactile domains, thus these sensory domains are the focus of this investigation. Over-responsivity manifests as extreme adverse or avoidant response to sensory stimulation, such as covering ears and running from the room in response to a vacuum cleaner, blender or automatic flushing toilet (auditory domain). In the tactile domain, sensory modulation difficulties manifest as refusal to wear clothing, particularly underwear, leading to significant household disruption and social challenge.

We seek to investigate the neurological underpinnings of SOR to determine if there is a unique neural signature that can be used as a biomarker for neural plasticity with intervention. This study focuses on tactile over-responsivity (TOR) and auditory over-responsivity (AOR) in a broad neurodevelopment cohort, taking a Research Domain Criteria (RDoC) inspired “sensory-first” approach [6]. The goal is to compare direct assessment and parent report measures of TOR and AOR in a pediatric cohort and to explore the neural architecture of SOR in children across categorical diagnoses.

### Characterizing Sensory Overresponsivity in Children with Neurodevelopmental Disorders

Many parent reports are used for assessing sensory modulation [7]–[14]. The Sensory Processing-Three Dimensions: Inventory quantifies sensory domains (vision, hearing, touch, and movement) by modulation, discrimination, as well as sensory-based motor challenges [4], [15]–[17]. The Sensory Sensitivity Questionnaire and the Sensory Experiences Questionnaire characterize sensory modulation specifically for children with ASD [18], [19]. The Sensory Profile (SP) has been validated cross-culturally and across clinical cohorts using sensory quadrant and section scoring methodology [10], [14], [20]–[22]. The Short Sensory Profile (SSP), derived from the SP, has been used to differentiate typically developing children from children with ASD [11], [13], [23], [24]. The SSP and other parent reports have made significant contributions to the research and clinical understanding of sensory dysfunction and are critical for a “trait-based” assessment. However, inclusion of a structured clinician administered assessment of sensory modulation has been lacking. Hence, this study seeks to advance the field of sensory assessment by comparing the auditory and tactile over-responsive items for children with neurodevelopmental disorders, using parent report, SSP, and direct assessment, Sensory Processing-Three Dimensions: Assessment (SP-3D:A) [4], [15]. The SP-3D:A is ideally suited for this task, as it includes characterizations of SOR in both auditory and tactile domains [4].

### Neural Architecture of Sensory Processing to Date

Prior work has begun to characterize the neural underpinnings of sensory processing differences in children with ASD, SPD, and ADHD [25]–[28]. Chang et al. (2016) reported robust alterations of posterior white matter microstructure in children with broadly defined SPD relative to typically developing children (TDC) [27]. This investigation found strong correlations between the fractional anisotropy (FA), a measure of microstructural integrity, with parent report and direct assessment measures of tactile and auditory discrimination across all children. However, direct assessment of sensory discrimination showed stronger and more continuous mapping to underlying white matter integrity than the parent report measures. Furthermore, in children with ASD, Pryweller et al. (2014) reported decreased FA in the inferior longitudinal fasciculus (ILF), which correlated directly with measures of TOR (defensiveness), suggesting atypical connectivity between the limbic system and multisensory integration regions [29]. This finding offers a preliminary explanation for the dysregulated emotional valence applied to non-noxious tactile stimuli. In this study, we hypothesize that direct assessment of AOR and TOR will show strong inter-test agreement with corresponding parental report behaviors in a NDD cohort; and, that sensory-first categorization using direct assessment of AOR will identify a more succinct subset of white matter tracts than previously identified using parent report.

## Methods

### Demographics

#### Experiment One: Direct Auditory and Tactile Over-Responsive Phenotyping

A total of 304 participants took part in experiment 1: 128 typically developing children (62 female; age 9.8 +/− 2.9) and 176 children with NDD (65 female; age 8.9 +/− 3.1). The NDD group was composed of 100 children with SPD (55 female; age 8.5 +/− 3.0) and 76 children with ASD (10 female; age 9.6 +/− 3.0). ASD cohort inclusion included a community diagnosis of ASD, a score of ≥15 on the Social Communication Questionnaire (SCQ) and/or a score of ≥25 on the Autism Quotient (AQ), and a confirmed ASD classification with the Autism Diagnostic Observation Schedule (ADOS) [30]–[32]. Participants in the SPD and TDC groups scored below cut-off criteria on the AQ or SCQ. Participants in the SPD cohort had an SPD designation from a community occupational therapist and/or a score in the “Definite Difference” range (<2% probability) in one or more of the SP section scores.

Participants in this Sensory Processing Disorders Consortium project were recruited from the University of California, San Francisco (UCSF) Sensory Neurodevelopment & Autism Program, the STAR Institute in Denver, Colorado, and the Icahn School of Medicine at Mount Sinai in New York. All parents provided written consent on behalf of their children, while children provided informed assent in accordance with each site’s institutional review board. For this retrospective study, not all children were administered all measures. All children (n=304) were included for establishing cut-off scores; children who had both direct assessment using the SP-3D:A and parent report using SSP were included in the phenotype comparison (n=235).

#### Experiment Two: Structural Neural Assessment of Auditory Over-Responsivity

For structural Diffusion Tensor Imaging (DTI) analysis, we included 39 boys from UCSF who successfully completed direct sensory assessment and neuroimaging assessment (ASD, n=13 (age 11 years +/− 2), SPD, n=8 (age 11 years +/− 1) and TDC, n=18 (age 12 years +/− 1)). This cohort has been previously described in Chang et al., 2016 [27]. Due to a small sample size in the tactile domain, we constrained the DTI analysis to the auditory domain.

## Measures

### Sensory Phenotyping Measures

#### Parent Report: Short Sensory Profile Questionnaire

The SSP includes 38-items in which parents rate how often their child shows a particular sensory behavior using a five-point Likert scale ranging from always (1) to never (5). Higher scores reflect more sensory-typical behavior. To align with the SP-3D:A, we inverted the scoring with never (1) and always (5). Thus, higher scores, on both parent report and direct assessment, will reflect greater SOR. The SSP has high internal reliability (.90-.95) and shows sensory differences in up to 90% of children and adults with ASD compared to controls [13], [24]. To achieve an SOR specific score for the auditory and tactile domains, we chose items reflecting SOR behaviors by clinical consensus (TT, EJM, SS, LJM, RC, LP) (see Table 1).

**Table 1:** Short Sensory Profile Items for Tactile and Auditory Over-Responsivity. Items below were identified by the clinical team as reflecting SOR behavior.

| SSP TOR | SSP AOR |
| --- | --- |
| 1. Expresses distress during grooming | 1. Responds negatively to unexpected or loud noises |
| 2. Avoids going barefoot, especially in sand or grass | 2. Holds hands over ears to protect ears from sound |
| 3. Reacts emotionally or aggressively to touch |  |
| 4. Withdraws from splashing water |  |
| 5. Has difficulty standing in line or close to other people |  |
| 6. Rubs or scratches out a spot that has been touched |  |

*SOR= Sensory Over-Responsivity; SSP= Short Sensory Profile; TOR= Tactile Over-Responsivity; AOR= Auditory Over-Responsivity*

#### Clinician Administered Assessment: Sensory Processing-Three Dimensions: Assessment

The SP-3D:A, a structured observational tool measuring behavioral response to specific sensory stimuli, includes probes that are administered by a STAR Institute trained, research reliable experimenter. The internal reliability is high (alpha=.94) [4]. Here, we included three auditory probes: “Find a picture”, during which participants cross out symbols with loud background noise; “Orchestra time”, in which participants play along with loud music using provided instruments; and “Sound and pictures”, where participants identify sounds such as a vacuum cleaner or dog barking. The tactile probes included: “Paint your arm”, during which participants paint their arm with a feather, a brush, and a rough sponge; “Goo”, in which participants remove 2 plastic animals from goo; and “Fishing”, requiring participants to retrieve plastic fish from a bucket of ice water. The following SOR behaviors during the game are given a score of 0 (not present) or 1 (observed): adverse response (0/1), discomfort, worries, and/or avoidance (0/1). For auditory over-responsive (SP3D:AOR) and tactile over-responsive (SP3D:TOR) composite scores, we summated the SOR behavior scores. Behaviors observed during, not prior to or between tasks, are included. Thus each composite, SP3D:AOR and SP3D:TOR, ranges from 0-6.

### DTI Acquisition

MR imaging was performed on a 3T Tim Trio scanner (Siemens, Erlangen, Germany) using a 12 channel head coil with an axial 3D magnetization prepared rapid acquisition gradient-echo T1-weighted sequence (TE = 2.98 ms, TR = 2300 ms, TI = 900 ms, flip angle of 90) with in-plane resolution of 1 × 1 mm on a 256 × 256 matrix and 160 1.0 mm contiguous partitions. Whole-brain diffusion imaging was performed with a multislice 2D single-shot twice-refocused spin echo-planar sequence with 64 diffusion-encoding directions, diffusion-weighting strength of b=2000 s/mm2, iPAT reduction factor of 2, TE/TR =109/8000 ms, NEX=1, interleaved 2.2 mm-thick axial slices with no gap, and in-plane resolution of 2.2 mm × 2.2 mm on a 100×100 matrix. An additional image volume was acquired with no diffusion weighting (b = 0s/mm2). The total diffusion acquisition time was 8.7min. Structural MRI for all children was reviewed by Dr. Pratik Mukherjee, a pediatric neuroradiologist, blind to cohort assignment. No clinically significant structural anomalies were identified.

### DTI Pre-processing

The diffusion-weighted images were corrected for motion and eddy currents using Functional Magnetic Resonance Imaging of the Brain Software Library’s linear Image Registration Tool (FSL; FLIRT1) with 12-parameter linear image registration [33]. All diffusion-weighted volumes were registered to the reference b = 0s/mm2 volume. To evaluate participant movement, we calculated a scalar parameter quantifying the transformation of each diffusion volume to the reference. The non-brain tissue was removed using the Brain Extraction Tool. FA was calculated using FSL’s DTIFIT at every voxel, yielding FA maps for each participant.

### Region of Interest DTI Analysis

Tract-Based Spatial Statistics in FSL was used to skeletonize and register the diffusion maps for each participant in order to perform voxel-wise comparisons along the white matter skeleton [34]. First, each participant’s FA map was non-linearly registered to each other participant’s FA map to identify the most representative FA map as a registration target. The registered maps were then averaged and skeletonized to the center of the white matter. Next, each participant’s FA data was projected onto this mean skeleton to obtain skeletonized FA maps per participant. Tract regions of interest (ROIs) were created according to The Johns Hopkins University ICBM-DTI-81 White-Matter Labeled Atlas [35]. Right and left hemisphere ROI tracts were highly correlated (r≥.50, p≤.001), thus an average diffusion value across right and left tracts were created for each participant.

## Statistical Analysis

### Experiment One: Cut score analysis and inter-test reliability

SPSS 24 was used to analyse the SSP and SP-3D:A data. Cut scores were designated at one standard deviation above the TDC group’s mean (rounded to the nearest integer). Inter-rater reliability was calculated by measuring the absolute agreement between SSP:AOR and SP3D:AOR, and between SSP:TOR and SP3D:TOR. Chi-square analysis was used to assess differences in over-responsivity between the NDD and TDC group.

### Experiment Two: DTI analysis between children with and without Auditory Over-Responsivity

Utilizing the SP3D:AOR cut-score determined in experiment one, we categorized the neuroimaging cohort to either an AOR (n=15) or NO-AOR (n=24) cohort. Due to a small sample size in the tactile domain (n=8), we focused on AOR for Experiment 2. We analyzed mean FA differences in 22 bilateral ROIs. We constructed ANOVAs using the categorical predictor variable for AOR (2 levels: above or below cut-score) and the outcome variables were the 22 ROIs. We review these findings both with and without False Detection Rate (FDR) correction to p-values (0.05) for each ANCOVA test.

## Results

### Experiment 1

Cut-off scores, 1 SD above the TDC mean, for SOR on parent report and direct assessment measures were established (see Table 2). Using direct assessment, children were classified as SP3D:AOR or SP3D:TOR if they scored 2 or above. With these direct assessment cut-off scores, 31% of children with NDD were classified as having AOR and 27% having TOR (Table 3). Using the SSP parent report, children were classified as SSP:AOR if they scored 5 or above and SSP:TOR if they scored 11 or above. Thus, using parent report, 62% of the children with NDD were classified as having AOR, whereas 68% had TOR. The Inter-test-agreement between SSP and SP-3D:A for AOR was 65% and TOR was 50%. Based on a two-proportion z-test for SP3D:AOR, SSP:AOR, SP3D:TOR, and SSP:TOR, the NDD group was significantly more affected by SOR than the TDC group (χ2≥17.5, p≤.0001).

**Table 2.** Cut-off scores for auditory and tactile over-responsivity. Cut scores were designated at 1 Standard Deviation (SD) above the TDC group’s mean (rounded to the nearest whole integer).

|  | TDC Mean<br>± SD (n) | Cut-Score | NO-SOR | SOR |
| --- | --- | --- | --- | --- |
| SP-3D:A - Auditory | .38 ± .98 (127) | 1.3 | 0-1 | ≥ 2 |
| SP-3D:A - Tactile | .29 ± .72 (128) | 1.0 | 0-1 | ≥ 2 |
| SSP - Auditory | 3.0 ± 1.4 (89) | 4.4 | 2-4 | ≥ 5 |
| SSP - Tactile | 7.5 ± 2.2 (92) | 9.7 | 6-10 | ≥ 11 |

*TDC= Typically Developing Children; SOR = Sensory Over-Responsivity; SP-3D:A = Sensory Processing- Three Dimensions:Assessment; SSP= Short Sensory Profile Parent Report*

**Table 3.**
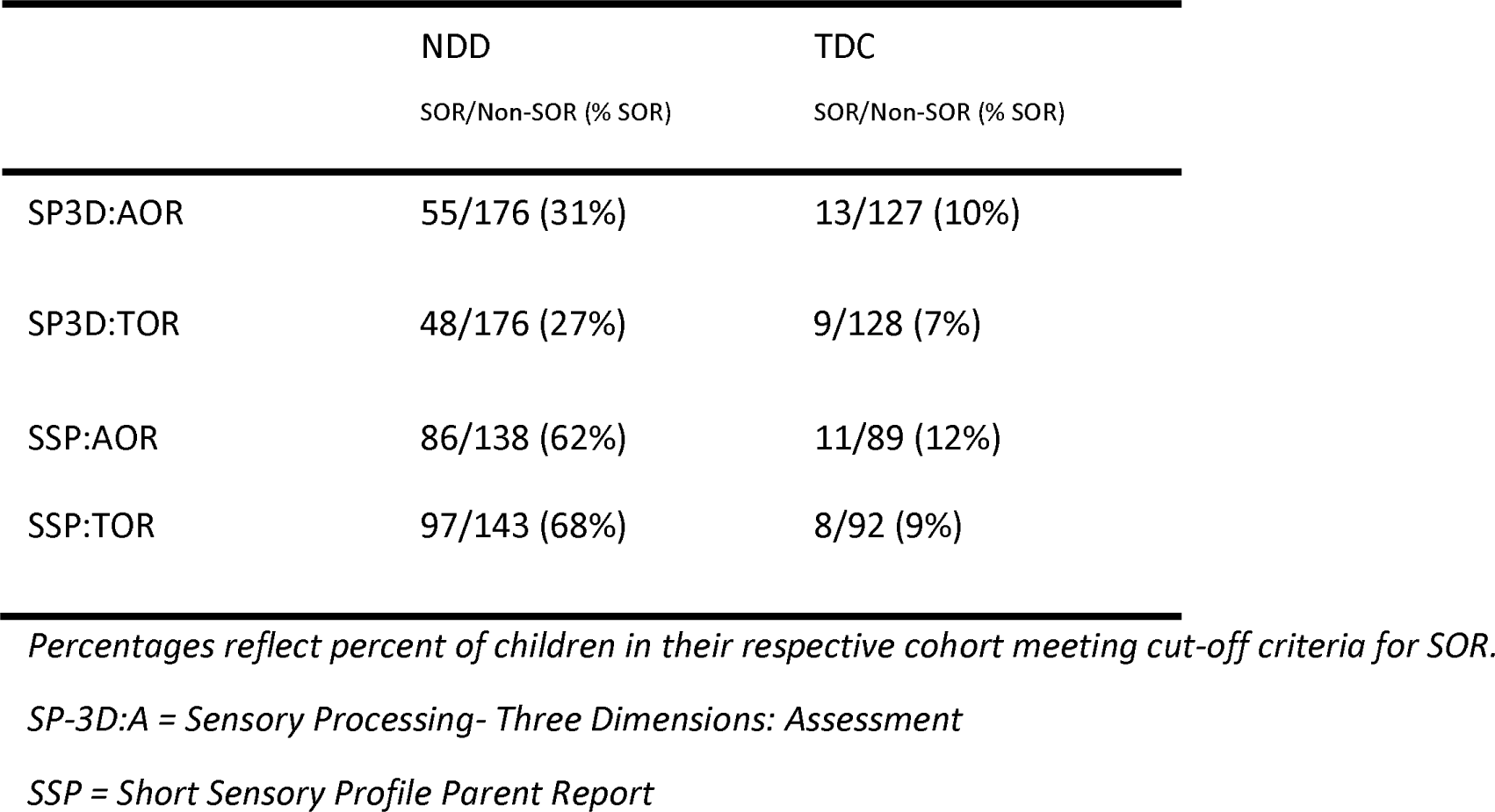
Count and Percentage of children with auditory or tactile over-responsivity

### Experiment 2

The second aim of our study was to identify the neural mechanisms contributing to AOR based on direct assessment. We compared DTI tracts from children who also completed the SP-3D:A. Based on our SP3D:AOR cut score analysis, 15 children (3 TDC, 7 ASD, 5 SPD) met AOR threshold and 24 did not. The AOR and NO-AOR cohorts did not differ in age (p=.37), perceptual-IQ (p=.35), or verbal-IQ (p=.53). We found that children with AOR had 11 total tracts showing decreased FA relative to children without AOR. Given the concern for multiple comparisons with this data driven approach, we applied FDR correction and 3 tracts continued to exceed the specified p value of < 0.05. These tracts are the posterior corona radiate (PCR), cingulate gyrus-cingulum portion (CGC), and superior longitudinal fasciculus (SLF; See Table 4 and Figure 1).

**Figure 1.**
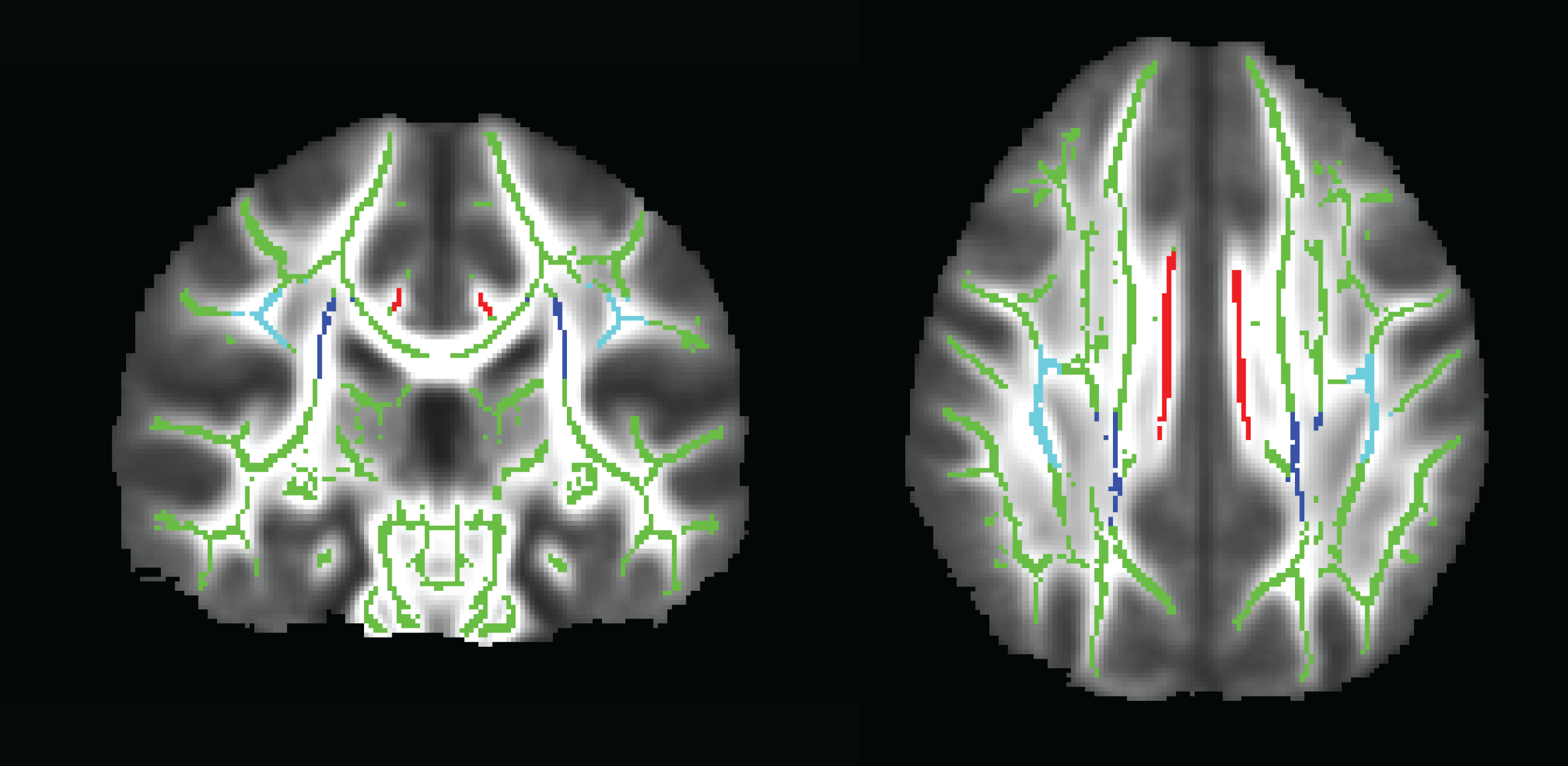
Skeletonized map of FA tracts. Image of the FA skeleton mask (green) displaying the tracts with significantly lower FA in the AOR group; the bilateral posterior corona radiata (PCR; dark blue), superior longitudinal fasciculus (SLF; light blue), and the cingulate gyrus-cingulum portion (CGC; red).

**Table 4.**
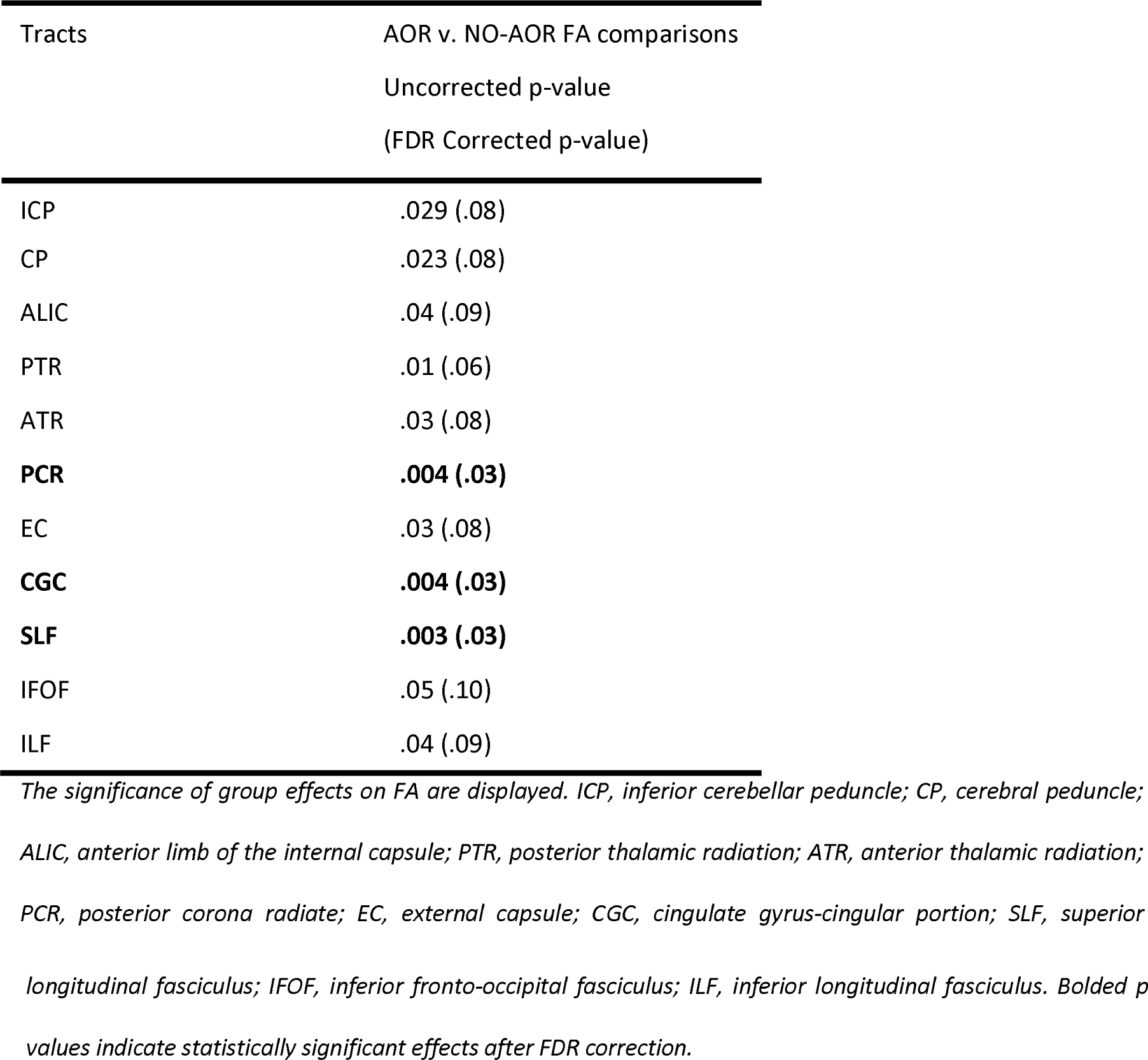
DTI tracts showing decreased FA in the auditory over-responsive cohort

## Discussion

Sensory processing dysfunction, specifically sensory over- and under-responsivity are now part of the DSM-5 criteria for ASD [5]. However, sensory processing challenges are also reported in children with other categorical conditions including ADHD, anxiety, and as the principle behavioral symptom for children with isolated Sensory Processing Disorder. This growing recognition has motivated the need for better clinical and research measures to characterize sensory modulation. Here, in line with the RDoC framework, we investigate SOR in the auditory and tactile domains as a dimension independent of clinical condition. We show that auditory and tactile over-responsivity can be quantified directly for children with and without NDD and that direct assessment has moderate concordance with parent report measures. Second, we report three neural tracts that differentiate children with AOR from those without.

Development of reliable sensory tools, both parent reports and direct assessments, is a critical step for researchers and clinicians alike. We hypothesized that AOR and TOR group assignment utilizing a combined parent report/direct assessment methodology, similar to that used as gold standard diagnosis in ASD, would provide a more robust sensory cohort assignment and that this combined assessment might be more robust for use with structural neuroimaging analysis. However, we found that the parent report questionnaire and direct observation have only a moderate overlap. Specifically, the agreement between SSP and SP-3D:A for AOR was 65% and TOR was 50%. This finding is similar to previous work showing moderate agreement between a sensory questionnaire and direct observation [15], [36]. Tavassoli et al. (2016) found an inter-rater agreement between questionnaire and observation of 74%, however, general sensory processing was evaluated rather than auditory and tactile over-responsivity [36]. Schoen et al (2008) focussed on SOR and reported a moderate correlation of .47, similar to our findings [15]. In line with previous reports, we find that more children meet SOR criteria based on parent report than on direct assessment in both the auditory and tactile domains, suggesting that the direct assessment may be a more stringent measure [36]. There are several plausible explanations; first, a direct observation might be more robust because of the subjective nature of questionnaires, parental bias, and recollection bias. Thus, using a direct assessment of the child’s observed behavior could allow for a more reliable and reproducible measure of sensory processing dysfunction. A second alternative explanation is that parents have more chances to observe their child’s sensory reactivity symptoms across various environments. In a laboratory setting, the amount of sensory stimuli is controlled for and does not represent the vast amount of sensory stimuli a child might experience in everyday situations. Therefore, parent reports likely reflect their child’s behavior across settings. For clinical utility, we suggest using a combination of measures to identify children at risk, such as a sensory questionnaire and clinical assessment. The goal is to detect all children who might have sensory modulation challenges that interfere with learning and social engagement and to be able to clinically intervene as early as possible. For research purposes, however, we suggest the use of sensory questionnaires as a screening tool, followed by standardized direct observations. Quantitative direct observation measures should be used when investigating biological mechanisms. However, future research with larger sample sizes and testing across multiple domains is needed to test these assumptions.

We previously reported significant differences in white matter microstructure in children with SPD and ASD relative to TDC [26]. However, as we have reported in our functional somatosensory magnetoencephalography work, neural mechanisms can often be better understood by splitting groups not by a clinical label, such as ASD, but by a more narrow construct of interest, such as tactile sensitive versus tactile typical [37]. Engaging a similar approach in this investigation, we split our cohort not by traditional clinical labels (ASD, SPD or TDC) but by a direct measure of AOR.

We conjectured that a sensory-first phenotype, in this case AOR, allows for a more parsimonious identification of key neural tracts. Indeed, in our previous work based on parent report and a broad inclusion criteria for sensory processing dysfunction, we found decreased FA in children with SPD in the posterior body and isthmus of the corpus callosum, the left posterior thalamic radiations (PTR), left PCR, and the posterior aspect of the left SLF [27]. Here, for children with AOR, the PCR, CGC, and SLF tracts showed decreased FA. In this analysis, the isthmus, posterior body of the corpus callosum, and the PTR were not significantly different between AOR and NO-AOR cohorts. While one might postulate that the current analysis was underpowered to detect the difference, this is unlikely given that the original study had 16 children in the general SPD group and 24 children in the TDC group, which is roughly similar to the 15 AOR and 24 NO-AOR children in this present study. We submit, instead, that the PCR, SLF, and CGC may represent critical connections in an AOR network. Additional work in a larger sample that will allow for investigation of TOR to determine if this network is a shared over-responsivity network or specific to the auditory domain is needed. In addition, a larger sample will allow for comparisons of SOR architecture in children with additional neurodevelopmental domains of challenges such as dysgraphia, dyspraxia, or sustained attention deficits.

With this study, we are also able to consider the concordance between tracts previously implicated in auditory discrimination with those implicated in auditory modulation. It is of note that all three of the tracts identified in this modulation-based approach, the PCR, CGC, and SLF, were also found to correlate with a global measure of non-linguistic auditory discrimination using the Differential Screening Test for Processing (DSTP). The DSTP showed a much broader correlation with anterior and posterior association tracts as well as commissural and projection tracts including the PTR. Thus, we posit that either the modulation network represents a subset of the discrimination networks, or that children included in this study are co-morbidly affected by both auditory discrimination and modulation dysfunction, which has been previously reported in children with ASD [4], [36], [38]. A larger and more heterogeneous neurodevelopmental sample will allow for these contrasts.

### Future directions and limitations

As with any study, there are limitations. Although over 300 participants took part in our first analysis, only 39 participants took part in the DTI imaging experiment. Consequently, the TOR cohort with neuroimaging available consisted of only 8 children which was insufficient for statistical comparison. For future SOR neuroimaging studies, a larger group of children with mixed neurodevelopmental profiles will allow for a broader range of sensory dysfunction. Furthermore, large and broad NDD cohorts will facilitate the understanding of whether SOR differences are fundamentally related to the current categorical cohorts such as ASD, ADHD, and SPD. However, emerging genetic findings, imaging reports, and even overlap in clinical semiology for individual children suggest that SOR will not respect these clinical divisions. Another limitation is that the Cingulum Bundle was divided into two portions, the superior and hippocampal region. While this is standard convention, reports that suggest a finer parcellation of the CGC into retrosplenial and subgenual divisions to better reflect the independent connections should be considered [39]. Finally, the concordance between sensory questionnaire and direct observation was only moderate. This was expected due to the different nature of measures and previous work [15], [36]. Future studies should determine if other parent-report measures have higher overlap with direct assessment.

## Conclusions

This study identified cut scores for AOR and TOR using parent report measure and direct observation. The SSP parent report and SP-3D:A direct observation ratings overlapped moderately for AOR and TOR. The direct observation measure here, the SP-3D:A, can be used in clinical and research settings to augment SOR phenotyping and investigate underlying mechanisms of sensory modulation and discrimination independently.

## List of Abbreviations

TOR: Tactile Over-Responsive
AOR: Auditory Over-Responsive
SSP: Short Sensory Profile
SP-3D:A: Sensory Processing-Three Dimensions: Assessment
SPD: Sensory Processing Dysfunction
NDD: Neurodevelopmental Disorders
ASD: Autism Spectrum Disorder
DSM-5: Diagnostic and Statistical Manual-5
SOR: Sensory Over-Responsive
RDoC: Research Domain Criteria
SP: Sensory Profile
TDC: Typically Developing Children
FA: Fractional Anisotropy
ILF: Inferior Longitudinal Fasciculus
SCQ: Social Communication Questionairre
AQ: Autism Quotient
ADOS: Autism Diagnostic Observation Schedule
DTI: Diffusion Tensor Imaging
ROI: Region of Interest
FDR: False Detection Rate

## Declarations

### Ethics approval and consent to participate

Study approval was obtained from the Institutional Review Board of the University of California, San Francisco (10-01940). Written consent was collected from parents of the participants.

### Consent for publication

All recruited participants/parents have given consent for publication during the recruitment process.

## Availability of data and material

The datasets used and/or analyzed during the current study are available from the corresponding author on reasonable request.

## Authors’ contributions

TT contributed to data collection, data analysis, and manuscript prepararion. RC, LP, SS, and LJM contributed to data collection, concept formation, and manuscript review. MRG contributed to manuscript preparation and submission. JO and PM contributed to data processing, data analysis, concept formation, and manuscript preparation. EJM contributed concept formation, data collection and analysis, and manuscript preparation. All authors have read and approved the manuscript.

## Acknowledgements

We would like to thank Carly Demopoulos and Sam Payabvash for editorial review. We would also like to thank all the children and families for their participation and support in our research.

## References

[1] C. E. Robertson and S. Baron-Cohen, “Sensory perception in autism,” Nat. Rev. Neurosci., vol. 18, no. 11, pp. 671–684, Sep. 2017.

[2] E. J. Marco, L. B. N. Hinkley, S. S. Hill, and S. S. Nagarajan, “Sensory processing in autism: a review of neurophysiologic findings.,” Pediatr. Res., vol. 69, no. 5 Pt 2, p. 48R–54R, May 2011.

[3] R. R. Ahn Miller, L. A., Milberger, S. & McIntosh, D. N., R. R. Ahn, L. J. Miller, S. Milberger, and D. N. McIntosh, “Prevalence of parents’ perceptions of sensory processing disorders among kindergarten children,” Am J Occup Ther, vol. 58, no. 3, pp. 287–293, 2004.

[4] S. A. Schoen, L. J. Miller, and J. C. Sullivan, “Measurement in Sensory Modulation: The Sensory Processing Scale Assessment,” Am. J. Occup. Ther., vol. 68, no. 5, pp. 522–530, 2014.

[5] American Psychiatric Association, “Diagnostic and Statistical Manual of Mental Disorders, 5th Edition (DSM-5),” Diagnostic Stat. Man. Ment. Disord. 4th Ed. TR., p. 280, 2013.

[6] B. N. Cuthbert and T. R. Insel, “Toward the future of psychiatric diagnosis: the seven pillars of RDoC.,” BMC Med., vol. 11, p. 126, Jan. 2013.

[7] G. T. Baranek, B. A. Boyd, M. D. Poe, F. J. David, and L. R. Watson, “Hyperresponsive sensory patterns in young children with autism, developmental delay, and typical development.,” Am. J. Ment. Retard., vol. 112, no. 4, pp. 233–45, Jul. 2007.

[8] P. P. P. Cheung and A. M. H. Siu, “A comparison of patterns of sensory processing in children with and without developmental disabilities,” Res. Dev. Disabil., vol. 30, no. 6, pp. 1468–1480, Nov. 2009.

[9] J. Ermer and W. Dunn, “The sensory profile: A Discriminant Analysis of Children With and Without Disabilities.,” Am. J. Occup. Ther., vol. 52, no. 4, pp. 283–290, 1998.

[10] M. A. Kientz and W. Dunn, “A comparison of the performance of children with and without autism on the sensory profile,” Occup. Ther. assessments, vol. 51, no. 7, pp. 530–537, 1997.

[11] A. E. Lane, S. J. Dennis, and M. E. Geraghty, “Brief report: Further evidence of sensory subtypes in autism,” J. Autism Dev. Disord., vol. 41, no. 6, pp. 826–831, Jun. 2011.

[12] A. E. Lane, R. L. Young, A. E. Z. Baker, and M. T. Angley, “Sensory processing subtypes in autism: Association with adaptive behavior,” J. Autism Dev. Disord., vol. 40, no. 1, pp. 112–122, Jan. 2010.

[13] S. D. Tomchek and W. Dunn, “Sensory processing in children with and without autism: a comparative study using the short sensory profile.,” Am. J. Occup. Ther., vol. 61, no. 2, pp. 190–200, 2007.

[14] L. D. Wiggins, D. L. Robins, R. Bakeman, and L. B. Adamson, “Breif report: Sensory abnormalities as distinguishing symptoms of autism spectrum disorders in young children,” J. Autism Dev. Disord., vol. 39, no. 7, pp. 1087–1091, Jul. 2009.

[15] S. A. Schoen, L. J. Miller, and K. E. Green, “Pilot study of the Sensory Over-Responsivity Scales: assessment and inventory.,” Am. J. Occup. Ther., vol. 62, no. 4, pp. 393–406.

[16] B. A. Boyd et al., “Sensory features and repetitive behaviors in children with autism and developmental delays,” Autism Res., vol. 3, no. 2, pp. 78–87, Apr. 2010.

[17] S. A. Schoen, L. J. Miller, and J. Sullivan, “The development and psychometric properties of the Sensory Processing Scale Inventory: A report measure of sensory modulation,” J. Intellect. Dev. Disabil., vol. 42, no. 1, pp. 12–21, 2017.

[18] N. J. Minshew and J. A. Hobson, “Sensory sensitivities and performance on sensory perceptual tasks in high-functioning individuals with autism,” J. Autism Dev. Disord., vol. 38, no. 8, pp. 1485–1498, Sep. 2008.

[19] G. T. Baranek, F. J. David, M. D. Poe, W. L. Stone, and L. R. Watson, “Sensory Experiences Questionnaire: Discriminating sensory features in young children with autism, developmental delays, and typical development,” J. Child Psychol. Psychiatry Allied Discip., vol. 47, no. 6, pp. 591–601, Jun. 2006.

[20] L. Crane, L. Goddard, and L. Pring, “Sensory processing in adults with autism spectrum disorders,” Autism, vol. 13, no. 3, pp. 215–228, May 2009.

[21] W. Dunn, B. S. Myles, and S. Orr, “Sensory processing issues associated with Asperger syndrome: A preliminary investigation,” Am. J. Occup. Ther., vol. 56, no. 1, pp. 97–102, 2002.

[22] R. L. Watling, J. Deitz, and O. White, “Comparison of Sensory Profile scores of young children with and without autism spectrum disorders.,” Am. J. Occup. Ther., vol. 55, no. 4, pp. 416–23, 2001.

[23] B. L. Brockevelt, R. Nissen, and E. Kurtz, “A Comparison of the Sensory Profile Scores of Children with Autism and an Age and Gender-Matched Sample,” Journal, vol. 66, no. November, pp. 459–465, Nov. 2013.

[24] D. N. McIntosh, L. J. Miller, V. Shyu, and V. Dunn, “Short Sensory Profile” in Sensory Profile user’s manual, 1999, pp. 59–73.

[25] J. P. Owen et al., “Abnormal white matter microstructure in children with sensory processing disorders.,” NeuroImage. Clin., vol. 2, pp. 844–53, Jan. 2013.

[26] Y. S. Chang et al., “Autism and sensory processing disorders: Shared white matter disruption in sensory pathways but divergent connectivity in social-emotional pathways,” PLoS One, vol. 9, 2014.

[27] Y.-S. Chang et al., “White Matter Microstructure is Associated with Auditory and Tactile Processing in Children with and without Sensory Processing Disorder,” Front. Neuroanat., vol. 9, Jan. 2016.

[28] S. A. Green et al., “Overreactive brain responses to sensory stimuli in youth with autism spectrum disorders.,” J. Am. Acad. Child Adolesc. Psychiatry, vol. 52, no. 11, pp. 1158–72, Nov. 2013.

[29] J. R. Pryweller et al., “White matter correlates of sensory processing in autism spectrum disorders.,” NeuroImage. Clin., vol. 6, pp. 379–87, Jan. 2014.

[30] C. Lord et al., “Autism diagnostic observation schedule: a standardized observation of communicative and social behavior.,” J. Autism Dev. Disord., vol. 19, no. 2, pp. 185–212, Jun. 1989.

[31] M. Rutter, A. Bailey, and C. Lord, SCQ: Social Communication Questionnaire. Los Angeles: Western Psychological Services, 2003.

[32] S. Baron-Cohen, S. Wheelwright, R. Skinner, J. Martin, and E. Clubley, “The Autism-Spectrum Quotient (AQ): Evidence from Asperger Syndrome/High-Functioning Autism, Malesand Females, Scientists and Mathematicians,” J. Autism Dev. Disord., vol. 31, no. 1, pp. 5–17, 2001.

[33] M. Jenkinson, P. Bannister, M. Brady, and S. Smith, “Improved optimization for the robust and accurate linear registration and motion correction of brain images,” Neuroimage, vol. 17, no. 2, pp. 825–841, Oct. 2002.

[34] S. M. Smith et al., “Tract-based spatial statistics: Voxelwise analysis of multi-subject diffusion data,” Neuroimage, vol. 31, no. 4, pp. 1487–1505, Jul. 2006.

[35] S. Wakana, H. Jiang, L. M. Nagae-Poetscher, P. C. M. van Zijl, and S. Mori, “Fiber Tract–based Atlas of Human White Matter Anatomy,” Radiology, vol. 230, no. 1, pp. 77–87, Jan. 2004.

[36] T. Tavassoli et al., “Measuring Sensory Reactivity in Autism Spectrum Disorder: Application and Simplification of a Clinician-Administered Sensory Observation Scale,” J. Autism Dev. Disord., vol. 46, no. 1, pp. 287–293, Jan. 2016.

[37] E. J. Marco et al., “Children with autism show reduced somatosensory response: an MEG study.,” Autism Res., vol. 5, no. 5, pp. 340–51, Oct. 2012.

[38] C. Demopoulos et al., “Shared and Divergent Auditory and Tactile Processing in Children with Autism and Children with Sensory Processing Dysfunction Relative to Typically Developing Peers,” J. Int. Neuropsychol. Soc., vol. 21, pp. 1–11, 2015.

[39] D. K. Jones, K. F. Christiansen, R. J. Chapman, and J. P. Aggleton, “Distinct subdivisions of the cingulum bundle revealed by diffusion MRI fibre tracking: Implications for neuropsychological investigations,” Neuropsychologia, vol. 51, no. 1, pp. 67–78, 2013.

